# The bHLH transcription factor MpHYPNOS regulates gemma dormancy in the liverwort *Marchantia polymorpha*

**DOI:** 10.1101/2022.04.25.488978

**Authors:** Hirotaka Kato, Nami Yoshimura, Mikako Yoshikawa, Hideyuki Matsuura, Kosaku Takahashi, Daisuke Takezawa, Tomoyuki Furuya, Yuki Kondo, Hidehiro Fukaki, Tetsuro Mimura, Kimitsune Ishizaki

## Abstract

- Dormancy is a key process employed by land plants to adapt to harsh terrestrial environments. The liverwort *Marchantia polymorpha* produces dormant propagules called gemmae for asexual reproduction. The plant hormone abscisic acid (ABA) plays important roles in regulating dormancy in both the seeds of flowering plants and the gemmae of *M. polymorpha*.
- Based on previous transcriptome analysis, we identified the basic helix-loop-helix transcription factor MpHYPNOS (MpHYP) as a key regulator of gemma dormancy.
- Knock-out mutants of *MpHYP* showed much higher germination rates of gemmae in gemma cups than ABA-related mutants, while the growth and development of these mutants resembled that of the wild type. Transient induction of *MpHYP* caused irreversible growth arrest of gemmae and thalli. Transcriptome and RT-qPCR analyses revealed that MpHYP represses the expression of cell cycle–related genes and induces ABA biosynthesis and ABA-responsive genes. Indeed, ABA levels increased in *MpHYP* overexpression lines and decreased in *Mphyp* knock-out lines. However, the growth arrest caused by *MpHYP* overexpression was not suppressed by a mutation in an ABA receptor gene.
- These findings suggest that MpHYP regulates gemma dormancy and thallus growth mainly through the ABA-independent pathway, providing clues about ABA-dependent and independent regulation of dormancy in land plants.

## Introduction

Dormancy is a key process that helps land plants adapt to harsh environmental conditions. Land plants produce stress-tolerant, dormant, reproductive units such as seeds and spores, whose growth is arrested until specific physiological and environmental signals are perceived. This delayed germination allows plants to withstand unfavorable conditions such as drought or low temperature and to disperse their progeny to distant locations via wind, water flow, or animals. In addition to sexual reproduction through spores, the liverwort *Marchantia polymorpha* propagates asexually by producing gemmae inside gemma cups that form on the dorsal side of the vegetative body/thallus. Gemmae are dormant inside gemma cups (Shimamura, 2016). Once gemmae fall out of the gemma cups due to a physical stimulus such as rain drops, they begin to germinate in the presence of light and water (Bowman, 2016). Gemma dormancy is regulated by several plant hormones, including abscisic acid (ABA), auxin, and ethylene (Eklund *et al*., 2015; Kato *et al*., 2015; Eklund *et al*., 2018; Li *et al*.,2020). ABA plays major roles in the initiation and maintenance of seed dormancy in flowering plants (Sano & Marion-Poll, 2021), and it was thus proposed that land plants share common mechanisms to regulate the dormancy of their progeny (Eklund *et al*., 2018). Auxin also plays a positive role in dormancy for both seeds and gemmae (Liu *et al*., 2013; Eklund *et al*., 2015). Mutants defective in auxin biosynthesis and signaling produce germinated gemmae inside of gemma cups (Eklund *et al*., 2015; Kato *et al*., 2017). Notably, ethylene plays opposite roles in regulating the dormancy of seeds vs. gemmae: While ethylene and its signaling factors promote the release of seed dormancy in dicot species [reviewed in (Longo *et al*., 2020)], ethylene treatment enhances the dormancy of gemmae in the gemma cup (Li *et al*., 2020).

Studies in flowering plants have revealed the basis of the ABA biosynthesis pathway and signaling mechanism [reviewed in (Sano & Marion-Poll, 2021)]. The first step of ABA biosynthesis involves the conversion of zeaxanthin to all-*trans*-violaxanthin by zeaxanthin epoxidase [ZEP; (Marin *et al*., 1996)], an enzyme encoded by *ABA1* in Arabidopsis *(Arabidopsis thaliana)*. ABA4 is important for the conversion of all-trans-violaxanthin to 9’-*cis*-violazanthin or 9’-*cis*-neoxanthin, although the enzymatic activity of ABA4 has not been demonstrated (Perreau *et al*., 2020). The first dedicated step in ABA biosynthesis is the production of xanthoxin by a 9-*cis*-epoxycarotenoid dioxygenase [NCED; (Schwartz *et al*.,1997)]. The resulting xanthoxin is then converted by a xanthoxin dehydrogenase (XD) into abscisic aldehyde (Gonzalez-Guzman *et al*., 2002), which is oxidized into ABA by an abscisic aldehyde oxidase [ABAO; (Seo *et al*., 2000)]. The ABA signal is perceived by the soluble ABA receptors PYRABACTIN RESISTANCE 1 (PYR1)/PYR1-LIKE (PYL)/REGULATORY COMPONENTS OF ABA RECEPTORS (RCARs) and inhibits the phosphatase activity of group A protein phosphatase 2C (PP2C-A) (Ma *et al*., 2009; Park *et al*., 2009). In the absence of ABA, PP2C-A directly inactivates subclass III Sucrose non-fermenting 1 (SNF1)-related protein kinase 2 (SnRK2) via dephosphorylation (Umezawa *et al*., 2009; Vlad *et al*., 2009). SnRK2s activate the ABA response by phosphorylating target proteins including the basic leucine zipper (bZIP) transcription factor ABA INSENSITIVE 5 [ABI5; (Nakashima *et al*., 2009)].

Phylogenomic studies have revealed that the components of ABA biosynthesis and signaling are largely conserved across land plants (Komatsu *et al*., 2020). *M. polymorpha* produces endogenous ABA (Li *et al*., 1994; Tougane *et al*., 2010) and contains single orthologs of all ABA biosynthetic genes except *XD* (Bowman *et al*., 2017). Exogenously supplied ABA causes growth inhibition, the accumulation of soluble sugars (such as sucrose), and enhanced desiccation tolerance, and it activates stress-related genes including *Late Embryogenesis Abundant-Like (LEAL)* genes in *M. polymorpha* (Tougane *et al*., 2010; Akter *et al*., 2014; Jahan *et al*., 2019). Such responses do not occur in loss-of-function mutants of *MpPYL1*, which encodes the major ABA receptor in vegetative tissues (Jahan *et al*.,2019). Constitutive expression of *PP2C-A* (Mp*ABI1*) and knockout of the downstream transcription factor gene *MpABI3A* also lead to ABA insensitivity and defects in gemma dormancy (Eklund *et al*., 2018).

Here, we demonstrate that the *M. polymorpha* basic helix-loop-helix (bHLH) transcription factor MpHYPNOS (MpHYP) plays a critical role in gemma dormancy. MpHYP belongs to the class VIIIb bHLH family, which includes Arabidopsis INDEHICHENT (IND) and HECATEs (HECs). IND and HEC play diverse roles in plants, including gynoecium development, but little is known about their roles in regulating dormancy. Transcriptome analysis and phytohormone quantification revealed that MpHYP promotes gemma dormancy partially via the regulation of ABA biosynthesis and ABA responses. Our study provides evolutionary insights into the roles of class VIIIb bHLH transcription factors and the cooperation of ABA-dependent and -independent dormancy regulation in land plants.

## Materials and Methods

### Plant materials, growth conditions, and transformation

Male and female accessions of wild-type *M. polymorpha*, Takaragaike-1 (Tak-1) and Tak-2 (Ishizaki *et al*., 2008), were maintained asexually. *M. polymorpha* plants were cultured as previously described (Yasui *et al*., 2019) unless otherwise indicated. Dexamethasone (DEX), cycloheximide (CHX), and ABA were dissolved in ethanol to prepare stock solutions and were added to agar medium after autoclaving. Agrobacterium *(Agrobacterium tumefaciens)-mediated* transformation of sporelings or thalli was performed as previously described (Ishizaki *et al*., 2008; Kubota *et al*., 2013).

### Generation of knock-out mutants of *MpHYP*

The 4.5- or 4.8-kb genomic region surrounding the region encoding the bHLH domain of MpHYP was amplified using the primer pairs MpHEC1_5IF_F and MpHEC1_5IF_R or MpHEC1_3IF_F and MpHEC1_3IF_R, and cloned into the *PacI* or *AscI* site of the pJHY-TMp1 vector, respectively (Ishizaki *et al*., 2013). The resulting plasmid was used to transform F1 sporelings generated by crossing Tak-2 and Tak-1. To confirm recombination, genomic PCR was performed using primer sets A (MpHEC1_GT_check_5_F and MpEF_GT_R1), B (H1F and MpHEC1_GT_check_3_R), and C (MpHEC1_5IF_check_F and MpHEC1_3IF_check_R). Sex determination was performed by genomic PCR with a mixture of four primers: rbm27_F, rbm27_R, rhf73_F, and rhf73_R. Genomic DNA was extracted as previously described (Hiwatashi *et al*., 2019).

### Complementation of *Mphyp^ko^*

A genomic fragment from 4.3 kb upstream of the transcriptional start site to the last codon of *MpHYP* was amplified using the primer set MpHEC1_pro_F and MpHEC1_CDS_nsR and cloned into the pENTR/D-TOPO vector (Thermo Fisher), followed by LR reaction with pMpGWB301 (Ishizaki *et al*., 2015) using LR Clonase II Enzyme Mix (Thermo Fisher). The downstream genomic region was then amplified using the primer set MpHEC1_CDS_IF_F and MpHEC1_3UTR_IF_R2 and integrated into the first binary plasmid with an In-Fusion HD Cloning Kit (Clontech) using the *BamHI* and *Asc*I sites, followed by transformation into *Mphyp^ko^* thalli.

### Overexpression of *MpHYP*

The coding sequence of *MpHYP* without the stop codon was amplified from Tak-1 cDNA using the primer set MpHEC1_CDS_F and MpHEC1_CDS_nsR and cloned into pENTR/D-TOPO (Thermo Fisher), followed by an LR reaction with pMpGWB313 (Ishizaki *et al*., 2015) to bring the Mp*HYP* sequence under the control of the *MpEF1α* promoter and in-frame with the sequence encoding the glucocorticoid receptor domain (GR). The resulting plasmid was used to transform Tak-1 thalli.

### Mutagenesis of *MpNCED*

Two oligo DNAs (NY009 and NY010) were annealed and cloned in pMpGE_En03 (Sugano *et al*., 2018). The resulting cassette expressing the sgRNA and Cas9 was transferred into the pMpGE010 vector using LR Clonase II Enzyme Mix (Thermo Fisher) and then transformed into Tak-1 thalli. To check the sequence at the Mp*NCED* locus, the genomic region around the sgRNA target site was amplified using the primer pair NY019 and NY020 and sequenced.

### Mutagenesis of the *MpPYL1* locus in *MpHYP-GR* plants

The plasmid expressing the sgRNA targeting *MpPYL1* and Cas9 (Jahan *et al*., 2019) was transformed into *MpHYP-GR* thalli. To check the sequence of the *MpPYL1* locus, the genomic region around the sgRNA target site was amplified using the primer set MpPYL1_qPCR_L1 and MpPYL1_qPCR_R1 and sequenced.

### Measuring germination rates of gemmae in gemma cups

Gemmae isolated from gemma cups located at the most basal positions in thalli were stained with 15 μM propidium iodide solution containing 0.01% (v/v) Triton X-100 for 15 min and observed under an M205 FA binocular fluorescence microscope (Leica) using the filter set for DsRED. The gemmae with elongated rhizoids were counted, and 79–114 gemmae were examined for each data point with three biological replicates.

### Scanning electron microscopy

For scanning electron microscopy (SEM), thalli were frozen in liquid nitrogen and observed under a VHX-D500 scanning electron microscope (KEYENCE).

### Promoter activity assay

A 6.3-kb genomic fragment including the 4.3-kb region upstream of the translational start site and the coding region before the conserved bHLH domain was amplified from Tak-1 genomic DNA using the primer pair MpHEC1_pro_F and MpHEC1_pro_R and cloned into the pENTR/D-TOPO vector. The promoter fragment was then transferred into pMpGWB304 (Ishizaki *et al*., 2015) to drive *GUS* expression. The resulting plasmid was introduced into Tak-1 thalli via Agrobacterium-mediated transformation (Kubota *et al*., 2013). Histochemical assays for GUS activity were performed as described previously with slight modifications (Ishizaki *et al*., 2012). The plant tissues were vacuum-infiltrated, followed by incubation in GUS assay solution for 4 h. To obtain section images, the samples were embedded in Technovit 7100 plastic resin as previously described (Yasui *et al*., 2019). Semi-thin sections (8-mm thick) were obtained with a microtome (HM 335E, Leica Microsystems, Germany) for light microscopy and stained with 0.01% (w/v) safranin O (Waldeck GmbH & Co. KG). The sections were observed under a BX51 upright microscope (Olympus) equipped with a DP74 CMOS camera (Olympus).

### Quantitative RT-PCR

Total RNA was extracted from the samples using an RNeasy Plant Mini Kit (Qiagen) and reverse transcribed into cDNA using ReverTra Ace qPCR RT Master Mix with gDNA remover (TOYOBO) according to the manufacturer’s protocol. Quantitative PCR was performed with a Light Cycler 96 instrument (Roche) using KOD SYBR qRT-PCR Mix (TOYOBO). The primers used in this study are listed in Table S1. The transcript levels of each gene were normalized to those of *MpEF1α* (Saint-Marcoux *et al*., 2015). Each measurement was performed with three or four biological replicates and three technical replicates.

### Transcriptome deep sequencing (RNA-seq)

Total RNA was isolated from the samples using an RNeasy Plant Mini Kit (Qiagen) following the manufacturer’s protocol. The extracted RNA was treated with an RNase-free DNase I set (Qiagen) and purified with an RNeasy MinElute Cleanup Kit (Qiagen). For RNA samples from WT and *Mphyp^ko^* gemma cups, sequencing libraries were constructed with a TruSeq RNA Sample Preparation Kit (Illumina) and sequenced on the HiSeq 4000 platform (Illumina) to obtain 100-bp paired-end data. For *MpHYP-GR* with or without 2 h of DEX treatment, sequencing libraries were constructed with a TruSeq Stranded mRNA Library Prep kit (Illumina) and sequenced on the NovaSeq 6000 platform (Illumina) to obtain 100-bp paired-end data. For *MpHYP-GR* with or without 24 h of DEX treatment, the sequencing libraries were constructed with a NEBNext Ultra II RNA Library Prep Kit for Illumina (NEB) and sequenced on the HiSeq 4000 platform to obtain 150-bp paired-end data. All resulting raw reads were deposited in the DDBJ Sequence Read Archive (DRA) under project accession number DRA013946.

Quality assessment of the raw RNA-seq reads was performed using FastQC (www.bioinformatics.babraham.ac.uk/projects/fastqc). Illumina adapters at the ends of the paired reads were removed using Cutadapt 2.1 (Martin, 2011). The clean reads were mapped onto the *Marchantia polymorpha* genome (v6.1 accessed through Marpolbase: https://marchantia.info/) using HISAT2 v2.1.0 (Kim *et al*., 2019) with default parameters. Post-processing of SAM/BAM files was performed using SAMTOOLS v1.9 (Li *et al*., 2009). GFF files were converted to GTF files using gffread embedded in cufflinks (Trapnell *et al*.,2010), and FeatureCounts v1.6.4 (Liao *et al*., 2014) was used to count the clean reads corresponding to each gene. EdgeR (Robinson *et al*., 2009) was used to normalize the raw counts and to perform differential gene expression analysis (*P*-adj < 0.01). Principal component analysis (PCA) was performed using normalized expression data for all genes. Gene Ontology (GO) enrichment analysis was performed using PlantRegMap by converting gene IDs from v6.1 to v3.1 (Tian *et al*., 2019).

### 5-Ethynyl-2’-deoxyuridine (EdU) incorporation assay

The Click-iT EdU Imaging Kit (Life Technologies) was used to visualize S-phase cells. EdU staining was performed as described previously (Furuya *et al*., 2021), with slight modifications. During sample clearing with ClearSee solution, cell walls were stained with SCRI Renaissance Stain 2200 (1:20,000 dilution). Alexa Fluor555 fluorescence was visualized under a FluoView FV1000 confocal laser-scanning microscope (Olympus) at excitation and detection wavelengths of 559 nm and 570–670 nm, respectively. Z-projection images were created using ImageJ software.

### Measurement of ABA contents

Plant tissue (ca. 1.0 g) was frozen in liquid nitrogen, crushed, and soaked overnight in 20 mL of ethanol. Ultra-performance liquid chromatography (UPLC) separation was performed using a Waters ACQUITY ethylene-bridged (BEH) C18 column (2.1-mm i.d. × 100 mm) and a Waters Micromass Quanttro Premier Tandem Quadrupole Mass Spectrometer (Waters, Milford, MA, USA). The endogenous ABA concentration was analyzed as described previously (Kobayashi *et al*., 2010).

## Results

### MpHYP plays a critical role in gemma dormancy

We previously compared transcriptome data obtained from whole thalli (TH), midribs (MR), and gemma cups (GCs) and identified 10 transcription factor genes that were highly expressed in GCs (Yasui *et al*., 2019). Among these, Mp5g18910 (Mapoly0073s0051 in the ver. 3.1 genome) showed the second highest expression levels in GCs relative to TH after *GEMMA CUP-ASSOCIATED MYB1 (GCAM1)*, encoding a critical regulator of gemma cup formation (Yasui *et al*., 2019). Confirmation by RT-qPCR revealed that Mp5g18910 shows an 11.5-fold higher expression level in GCs than in TH (Fig. 1a). Mp5g18910 comprised a single exon (Fig. 1b), and the encoded protein was phylogenetically classified within the class VIIIb bHLH subfamily, which includes Arabidopsis HECs and IND (Bowman *et al*., 2017; Bonnot *et al*., 2019). To analyze the biological function of Mp5g18910, we generated knock-out mutants by the homologous recombination method (Fig. 1b and Fig. S1; Ishizaki *et al*., 2013). We obtained two independent lines (#173 and #182) showing the same phenotype, with no obvious defects except for germinated gemmae inside gemma cups (Fig. 1c, d; Fig. S1). Wild-type (WT) Tak-1 gemmae were completely dormant inside gemma cups on 17-day-old thalli, and we observed no elongated rhizoids. By contrast, 27.8% of *Mphyp^ko^* mutant gemmae had already germinated inside gemma cups at the same stage (Fig. 1d,e). This dormancy defect was completely rescued by introducing the genomic fragment of Mp5g18910 (Fig. 1e). We thus named this gene *MpHYPNOS* (Mp*HYP*), after the personification of sleep in Greek mythology.

**Fig. 1.**
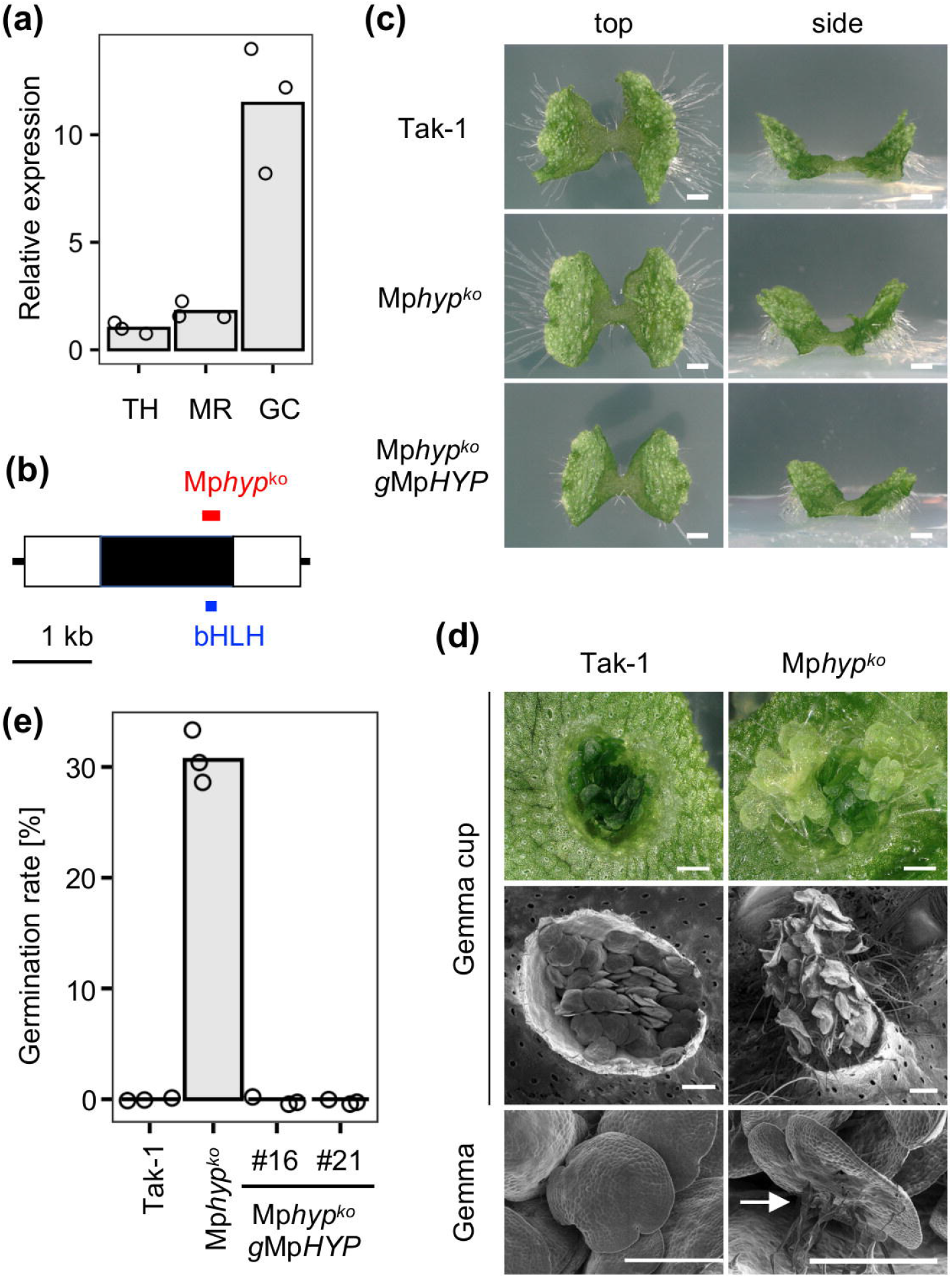
*MpHYP* plays a critical role in gemma dormancy. (a) Relative expression levels of Mp*HYP*, normalized to *MpEF1α*. Samples were collected from 1-week-old whole thalli (TH), midribs (MR), or gemma cups (GC) from 3-week-old thalli. Expression levels in TH were set to 1. Bars indicate mean values, and circles represent each data point. (b) Gene structure of Mp*HYP*. White and black boxes represent the untranslated regions (UTRs) and coding sequence, respectively. *MpHYP* does not contain introns. The blue bar indicates the region encoding the bHLH domain, and the red bar indicates the recombined region in the *Mphyp^ko^* mutants (see also Fig. S1a). (c) Top and side views of 1-week-old WT (Tak-1), *Mphyp^ko^*, and *Mphyp^ko^ gMpHYP* gemmalings. Bars = 1 mm. (d) Gemma cups and gemmae from 4-week-old WT and *Mphyp^ko^* thalli. Gray images were taken by SEM. The arrow indicates elongated rhizoids. Bars = 0.5 mm. (e) Germination rates of gemmae from 17-day-old WT (Tak-1), *Mphyp^ko^*, and *Mphyp^ko^ gMpHYP* plants. Bars indicate mean values, and dots represent each data point.

### *MpHYP* is expressed in the apical notch, midrib, and developing gemma

To identify the precise site of MpHYP function, we introduced a fusion construct harboring the Mp*HYP* promoter region driving the *β-glucuronidase (GUS)* reporter gene into WT gemmalings (Mp*HYP_pro_:GUS)*. We detected relatively weak GUS activity around the meristematic notches of young (5-day-old) gemmalings (Fig. 2a). At 10 days or at later stages, we observed strong GUS activity around meristematic notches, the apical region of the midrib, and GCs (Fig. 2b,c). Inside GCs, we obtained relatively high GUS activity in early developing gemmae and in the ventral side of the thallus (Fig. 2d,e). Interestingly, we observed little GUS activity in mature gemmae (Fig. 2f). These results suggest that *MpHYP* is expressed in the apical notch, midrib, GC, and developing gemma, which is consistent with the RNA-seq and RT-qPCR results (Fig. 1).

**Fig. 2.**
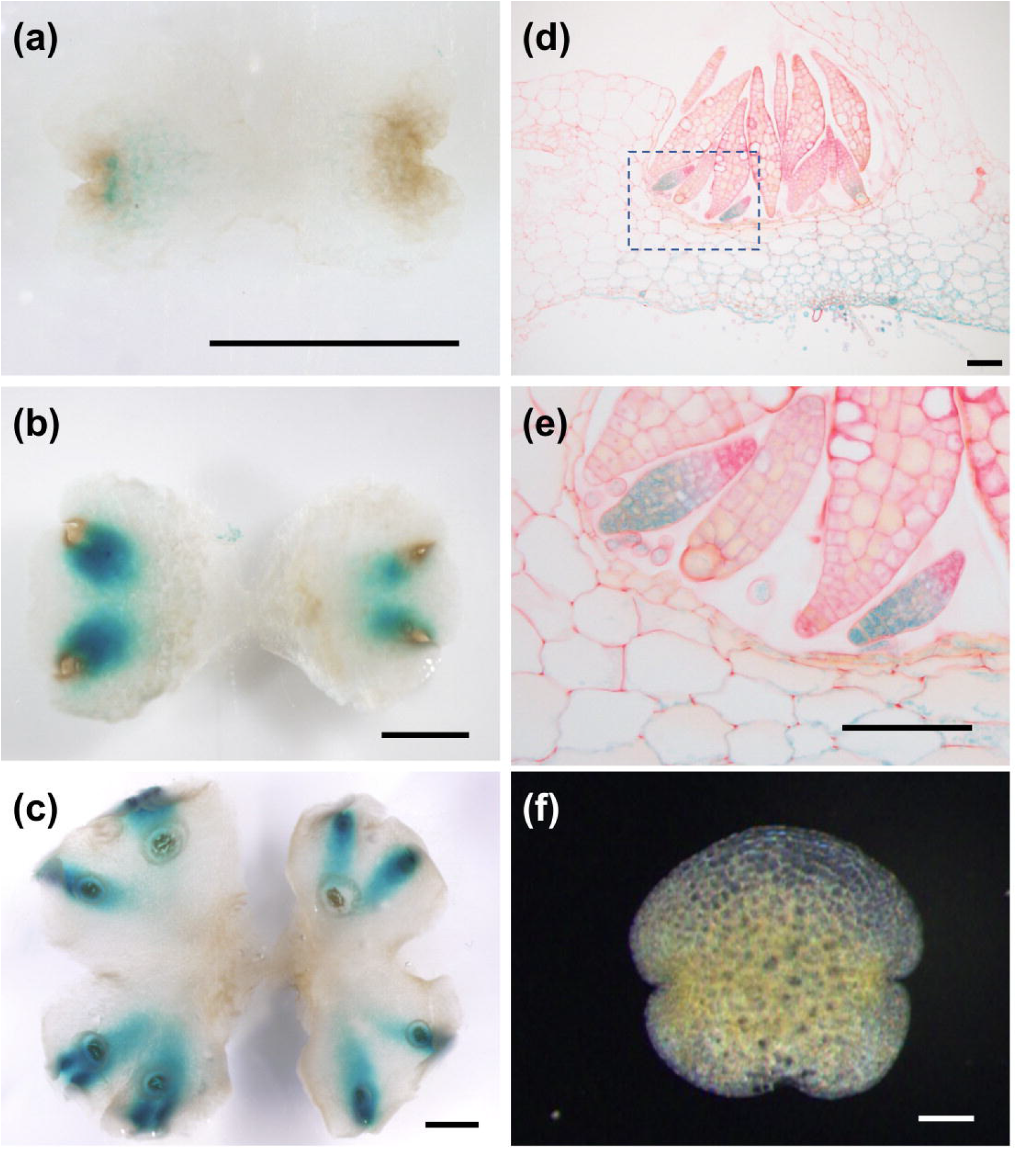
GUS staining of *MpHYP_pro_:GUS*. (a-c) Five-(a), 10-(b), and 14-day-old (c) *MpHYP_pro_:GUS* gemmalings. (d, e) Section images of gemma cups. Black squares indicate the regions magnified in (e). (f) Mature gemma before germination. Bars = 2 mm (a-c), 0.1 mm (d-f).

### Overexpressing *MpHYP* represses gemma germination and thallus growth

To investigate the effect of *MpHYP* overexpression, we generated transgenic plants producing a fusion protein between MpHYP and the glucocorticoid receptor domain (GR), whose expression was driven by the strong *MpEF1α (Elongation factor 1α)* promoter (Mp*HYP-GR*). MpHYP-GR protein should be active only in the presence of dexamethasone (DEX; Schena *et al*., 1991). The expression levels of Mp*HYP* in *MpHYP-GR* plants were more than 100-fold that of the WT (Fig. 3a). When *MpHYP-GR* plants were grown on DEX-containing medium, both ungerminated gemmae and 1-week-old thalli showed severe growth arrest (Fig. 3b,c). Interestingly, when the DEX-treated gemma or thalli were transferred to control medium, DEX-treated plants still exhibited severe growth arrest for at least 1 week (Fig. 3b,c). These results suggest that MpHYP strongly represses thallus growth as well as gemma germination in an irreversible manner.

**Fig. 3.**
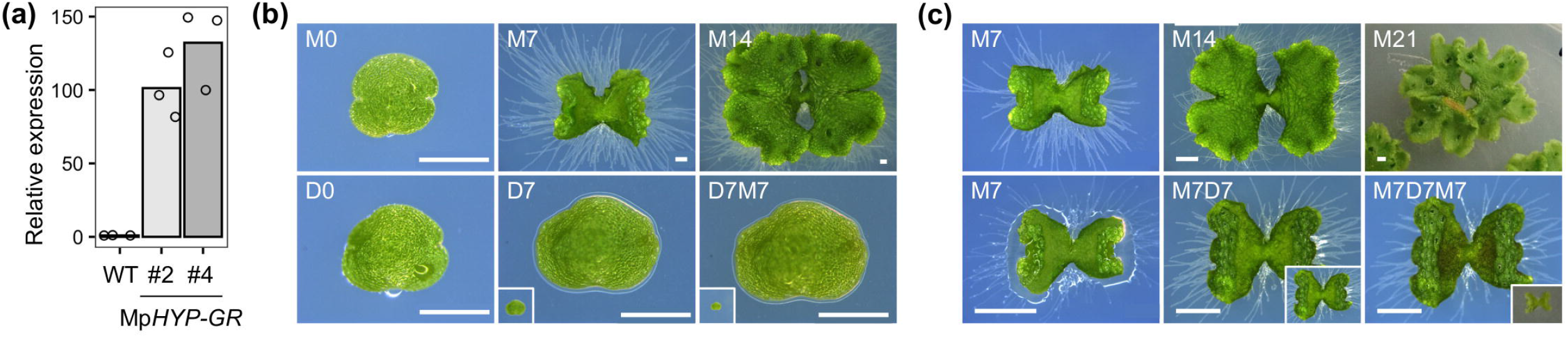
Overexpression of *MpHYP* represses gemma germination and thallus growth. (a) Expression analysis of *MpHYP* in Tak-1 (WT) and *MpHYP-GR* plants by RT-qPCR. RNA samples were collected from 1-week-old gemmalings. Expression levels were normalized to that of the WT. Bars indicate mean values, and dots represent each data point. (b, c) Gemmae (b) or 7-day-old gemmalings (c) of *MpHYP-GR* were grown in mock (M) or 10 μM DEX (D) conditions. The number indicates the days of each treatment. For example, M7D7 means that plants were grown for 7 days under mock conditions, followed by 7 days on medium containing DEX. Insets show images at the same magnification as tissues of the same age under mock conditions. Scale bars = 0.5 mm (b), 2 mm (c).

### MpHYP represses cell division activity

To explore the downstream pathway of MpHYP, we performed RNA-seq experiments using WT and *Mphyp^ko^* gemma cups and 1-week-old *MpHYP-GR* gemmalings following 2 or 24 h of DEX treatment. In the resulting principal component (PC) analysis plot, each biological replicate grouped close together, and both PC1 and PC2 values correlated with the direction of *MpHYP* genetic manipulation (Fig. 4a). Compared to WT gemma cups, we identified 2,750 upregulated genes and 2,222 downregulated genes in *Mphyp^ko^* gemma cups (Fig. 4b). When we compared 1-week-old *MpHYP-GR* gemmalings incubated with or without DEX, we obtained 2,700 and 4,250 upregulated genes, and 1,854 and 4,244 downregulated genes after 2 or 24 h of induction with DEX, respectively (Fig. 4b). Of these genes, 1,535 (57%) of upregulated genes and 1,316 (71%) of downregulated genes following 2 h of DEX treatment were also up- or downregulated after 24 h of this treatment (Fig. 4b). When we examined the overlap of differentially expressed genes (DEGs) across the three comparisons, we identified 412 downregulated genes in *Mphyp^ko^* and upregulated in *MpHYP-GR* at both time points, with 413 genes showing the opposite pattern (Fig. 4b). We defined these DEGs as MpHYP-activated and MpHYP-repressed genes, respectively.

**Fig. 4.**
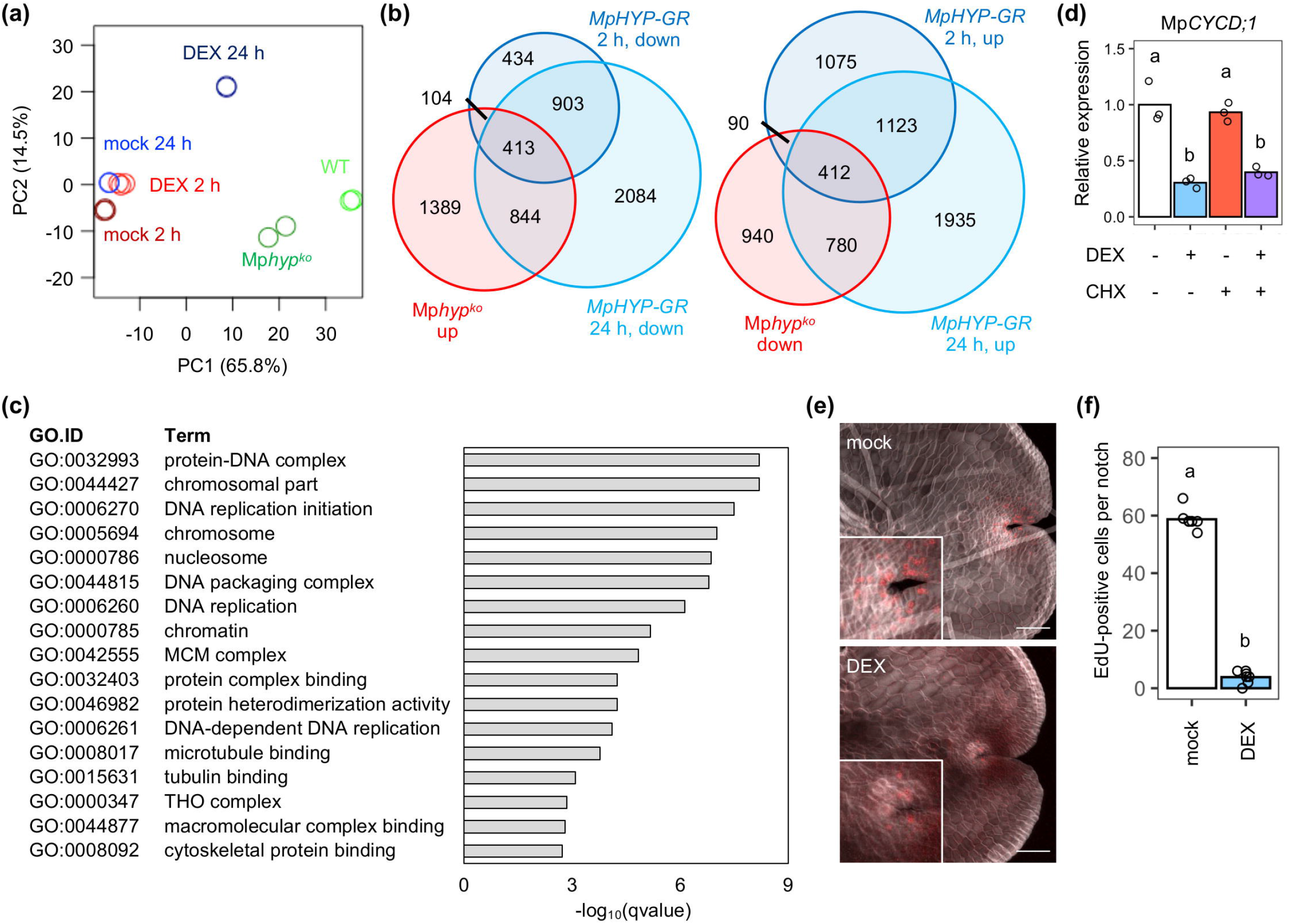
Transcriptome analysis using *Mphyp^ko^* and *MpHYP-GR*. (a) Principle component (PC) analysis plot of RNA-seq data from WT and *Mphyp^ko^* gemma cups and 7-day-old *MpHYP-GR* gemmalings treated without (mock) or with DEX for 2 or 24 h. Circles of the same color indicate biological replicates for the same conditions. (b) Venn diagrams of DEGs for each comparison. The number of genes classified in each group is indicated. (c) GO term enrichment analysis using the 413 MpHYP-repressed genes. (d) Expression analysis of *MpCYCD;1* by RT-qPCR. Seven-day-old *MpHYP-GR* gemmalings were treated with or without 10 μM DEX and/or 10 μM CHX for 2 h. Bars indicate mean values, and circles represent each data point (n = 3). Different lowercase letters indicate significant differences (*P* < 0.01 by ANOVA followed by Tukey’s HSD test). (e) EdU incorporation assay to visualize S phase progression in *MpHYP-GR*. Three-day-old gemmalings were incubated without (mock) or with 10 μM DEX for 6 h and cultured on medium containing EdU for 1 h. Images of EdU (red) and the cell wall (gray) were merged. Scale bars = 100 μm. (f) EdU-positive cells per notch were counted from the confocal microscopy images (e). Bars indicate mean values, and circles represent each data point (n = 7). Different lowercase letters indicate significant differences (*P* < 0.01 by ANOVA followed by Tukey’s HSD test).

We subjected the MpHYP-activated and MpHYP-repressed genes to Gene Ontology (GO) enrichment analysis to identify the biological pathways regulated by MpHYP. While no GO terms were significantly enriched (q-value < 0.01) among MpHYP-activated genes, many GO-terms related to DNA replication were enriched among MpHYP-repressed genes (Fig. 4c). Of the known cell cycle–related genes (Bowman *et al*., 2017), we noticed four genes encoding cyclins or cyclin-dependent kinase, which are important for cell cycle progression (Komaki *et al*., 2012), among MpHYP-repressed genes (Table S2). In particular, the expression of *MpCYCD;1*, encoding one of two D-type cyclins, which promote the transition from G1 to S phase in angiosperms and mosses (Masubelele *et al*., 2005; Ishikawa *et al*.,2011), decreased to approximately 30% within 2 h of the induction of *MpHYP-GR* by DEX treatment (Fig. 4d, Table S2). The repression of *MpCYCD;1* by *MpHYP-GR* was not suppressed by the translation inhibitor cycloheximide (CHX), suggesting that *MpCYCD;1* is a direct target of MpHYP (Fig. 4d). To determine whether overexpression of *MpHYP* represses DNA replication, we performed a 5-ethynyl-2’-deoxyuridine (EdU) incorporation assay, which visualizes the entry into S phase. While mock-treated gemmalings showed ~50 EdU-positive cells per apical notch, DEX-treated gemmalings had few or no EdU-positive cells (Fig. 4e,f). These results suggest that MpHYP represses cell division activity by repressing the expression of cell cycle–related genes.

### MpHYP promotes dormancy only partially through the ABA pathway

The plant hormones ABA, auxin, and ethylene positively regulate gemma dormancy in *M. polymorpha* (Eklund *et al*., 2015; Kato *et al*., 2017; Eklund *et al*., 2018; Li *et al*., 2020). To investigate whether these phytohormones participate in MpHYP-dependent dormancy regulation, we measured the expression levels of genes known to be involved in each phytohormone pathway in our RNA-seq data. No ethylene-related genes were found among either MpHYP-activated or -repressed genes (Table S3). For auxin-related genes, we identified *GRETCHEN HAGEN 3A (MpGH3A:* Mp6g07600), encoding an auxin-inactivating enzyme, among MpHYP-repressed genes (Table S4). However, no known auxin-activated genes appeared to be activated by MpHYP. These results suggest that MpHYP does not function via the regulation of the auxin or ethylene pathway.

We then focused on ABA-related genes, which revealed that a putative ABA biosynthesis gene, *MpABA4* (Mp6g13390), and 18 *MpLEAL* genes are activated by MpHYP (Tables S5 and S6). Twelve other *MpLEAL* genes and two other known ABA-responsive/signaling genes, *MpABI3A* and *MpABI5B* (Eklund *et al*., 2018), were significantly upregulated in *MpHYP-GR* at both time points, although the downregulation of these genes in *Mphyp^ko^* was not significant (Table S5). Notably, the ABA biosynthesis gene *MpNCED* (Mp2g07800) was rapidly and strongly upregulated in *MpHYP-GR* in the presence of DEX, which we confirmed by time-course RT-qPCR analysis (Fig. 5a; Table S5). By contrast, *MpABA4, MpLEAL1*, and *MpLEAL5* were upregulated only 4 h or later after the induction of *MpHYP-GR* by DEX (Fig. 5a). The activation of Mp*NCED* by DEX was not inhibited by the translation inhibitor CHX, while that of Mp*LEAL5* was (Fig. 5b). These results suggest that MpHYP directly activates Mp*NCED* expression, which results in increased ABA levels and transcriptional responses. To confirm this hypothesis, we measured the changes in ABA levels in response to *MpHYP* genetic modification. *Mphyp^ko^* accumulated lower levels of ABA in gemma cups than the WT, while DEX treatment increased ABA levels in *MpHYP-GR* thalli (Fig. 5c, d).

**Fig. 5.**
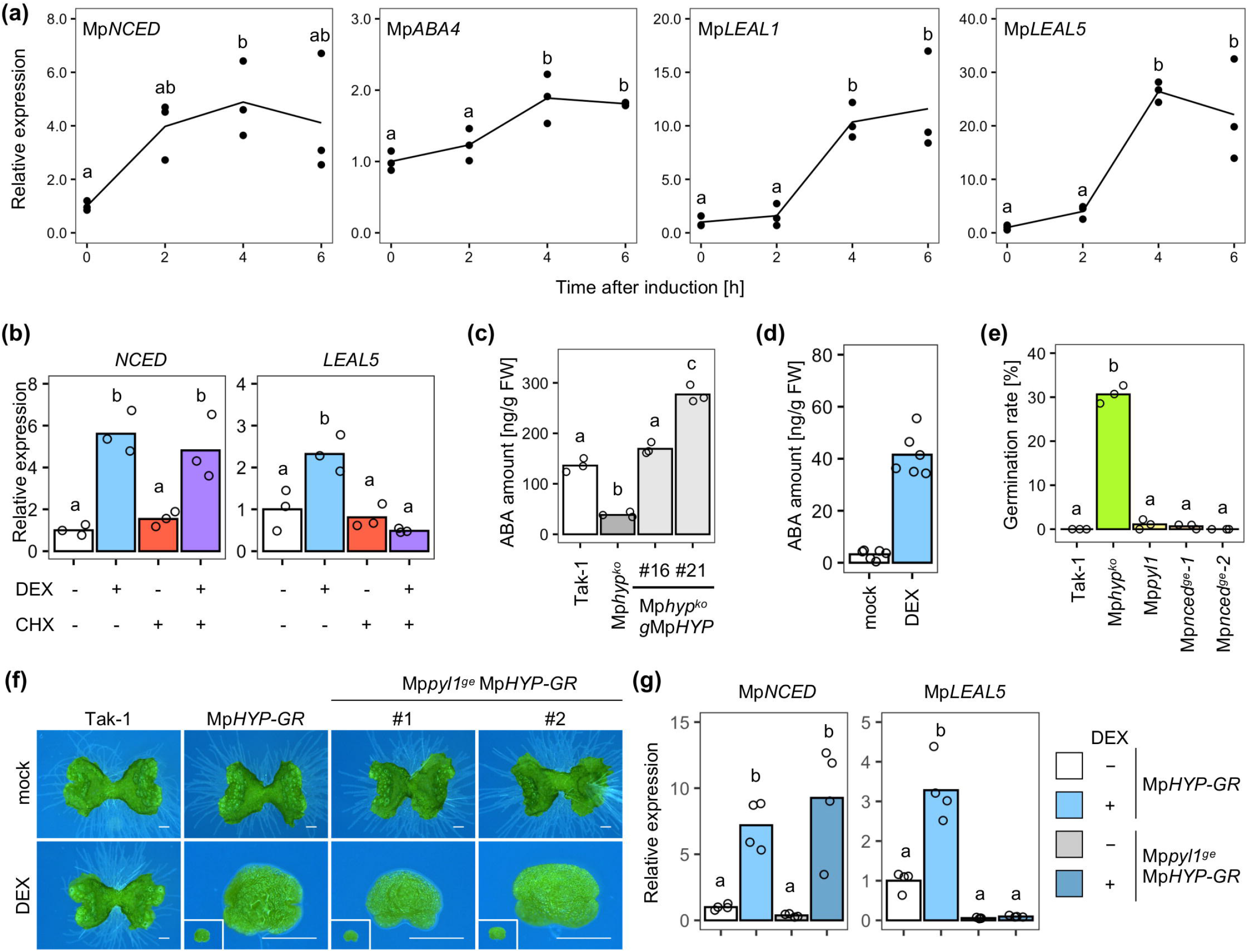
The relationship between *MpHYP* and the ABA pathway. (a) Time-course RT-qPCR analysis of *MpNCED, MpABA4, MpLAEL1*, and *MpLAEL5* expression in *MpHYP-GR*. Seven-day-old gemmalings were treated with 10 μM DEX for 0, 2, 4, or 6 h. Expression levels were normalized to the value at 0 h. Lines indicate mean values, and dots represent each data point. (b) RT-qPCR analysis of Mp*NCED* and *MpLEAL5* expression in 7-day-old *MpHYP-GR* thalli with or without 10 μM DEX and/or 10 μM CHX treatment for 2 h. Different lowercase letters indicate significant differences (*P* < 0.05 by ANOVA followed by Tukey’s HSD test, n = 3). (c, d) Quantification of ABA contents in WT, Mp*hyp^ko^*, and *Mphyp^ko^ gMpHYP* gemmae (c) and 1-week-old *MpHYP-GR* thalli with or without 10 μM DEX treatment for 24 h (d). (e) Germination rates of gemmae in gemma cups of WT, *Mphyp^ko^, Mppyl1*, and *Mpnced* at the 17-day-old gemmaling stage. (f) Seven-day-old gemmalings of Tak-1, *MpHYP-GR*, and *Mppyl1^ge^ MpHYP-GR* grown without or with 10 μM DEX. Bars indicate mean values, and circles represent each data point (a-e). Different lowercase letters indicate significant differences (*P* < 0.05, [a, b, g], or 0.01 [c-f] by ANOVA followed by Tukey’s HSD test).

To investigate whether an ABA-dependent pathway plays a major role in Mp*HYP*-dependent gemma dormancy, we compared the germination rates of gemmae in the WT, *Mphyp^ko^*, and the ABA receptor mutant *Mppyl1^ge2b^* (Jahan *et al*., 2019). In addition, we generated mutants for Mp*NCED* using clustered regularly interspaced short palindromic repeats (CRISPR)/CRISPR-associated nuclease 9 (Cas9)-mediated gene editing *(Mpnced-1^ge^* and *Mpnced-2^ge^;* Fig. S2a; Sugano *et al*., 2018). These mutants contained frameshift mutations in the beginning of the conserved region and thus should be loss-of-function alleles. Approximately 30% of *Mphyp^ko^* mutant gemmae had already germinated inside gemma cups of 17-day-old thalli, whereas most WT, *Mpyl1^ge2b^*, and *Mpnced* gemmae were still dormant at the same stage (Fig. 5e). Under our experimental conditions, we observed significant differences between the WT and *Mppyl1^ge2b^* or *Mpnced^ko^* only at a much later stage (28-day-old thalli; Fig. S2b).

We then generated loss-of-function mutants of *MpPYL1* in the *MpHYP-GR* background via CRISPR/Cas9-mediated gene editing *(Mppyl^ge^ MpHYP-GR* #1 and #2; Fig. S3a). *Mppyl^ge^ MpHYP-GR* plants were insensitive to exogenously supplied ABA, as were the *Mppyl1^ge2b^* mutants in the WT background (Fig. S3b; Jahan *et al*., 2019). Growth arrest caused by DEX treatment was not suppressed by the mutation of *MpPYL1* (Fig. 5f), suggesting that the contribution of the MpPYL1-dependent pathway to gemma dormancy or growth arrest caused by MpHYP is limited. Indeed, several ABA-independent pathways regulate ABA responses by controlling SnRK2 activity (Saruhashi *et al*., 2015; Née *et al*.,2017; Nishimura *et al*., 2018). To investigate whether similar bypass mechanisms might be present downstream of MpHYP, we measured the expression levels of ABA-responsive genes in *Mppyl1^ge^ MpHYP-GR* plants. While the expression of *MpNCED* was activated by DEX treatment regardless of the presence of MpPYL1, *MpLEAL5* expression was not induced by DEX in *Mppyl1^ge^ MpHYP-GR* plants (Fig. 5g). These results suggest that the activation of ABA-responsive genes by MpHYP is dependent on MpPYL1.

## Discussion

### MpHYP is a key regulator of gemma dormancy

Dormancy is a key process that allows plants to survive harsh environments on land. The liverwort *M. polymorpha* produces gemmae for asexual reproduction, which are dormant in the absence of germination signals such as light and water. Previous studies have indicated that the dormancy of gemmae, like seed dormancy in angiosperms, is regulated by ABA (Eklund *et al*.,2018). Based on our previous transcriptome analysis (Yasui *et al*., 2019), we identified the bHLH-type transcription factor MpHYP as a critical regulator of gemma dormancy in *M. polymorpha*. MpHYP directly promotes the expression of the ABA biosynthesis gene *MpNCED*, which results in increased ABA contents and the upregulation of ABA-responsive genes (Fig. 5, Tables S5-6). However, the finding that 1) *Mpnced^ge^* and *Mppyl1^ge^* mutants have lower gemma germination rates than *Mphyp^ko^* and 2) the same growth arrest phenotype caused by *MpHYP-GR* was observed regardless of the mutation of *MpPYL1* suggests that ABA biosynthesis and perception only partially contribute to MpHYP-dependent dormancy regulation. A previous study demonstrated that the ABA signal in gemmae, and not gemma cups, is important for maintaining gemma dormancy (Eklund *et al*., 2018). Here, histochemical analysis demonstrated that the *MpHYP* promoter is active in gemma cups and developing gemmae, but not in mature gemmae (Fig. 2). These data suggest that MpHYP is important for the establishment rather than the maintenance of gemma dormancy during gemma development, which might reflect the limited contribution of the ABA pathway.

Besides ABA, auxin and ethylene are known to regulate the dormancy of both *M. polymorpha* gemmae and the seeds of angiosperms (Liu *et al*., 2013; Corbineau *et al*., 2014; Eklund *et al*., 2015; Kato *et al*., 2017; Li *et al*., 2020). However, our transcriptome analysis suggested that MpHYP is not likely to function through these pathways. One candidate pathway for MpHYP-dependent dormancy regulation is the cell cycle. This study showed that MpHYP inhibits the expression of various cell cycle–related genes and entry into S phase (Fig. 4, Table S2). In particular, RT-qPCR of plants under CHX treatment suggested that MpHYP directly downregulates *MpCYCD;1*, whose expression was not inhibited by ABA in a previous RNA-seq analysis (Jahan *et al*., 2019). Interestingly, growth arrest caused by the transient activation of MpHYP-GR continued for at least 1 week after the DEX treatment was suspended, and this growth inhibition was independent of the ABA pathway (Fig. 3). Such long-term effects might be achieved through epigenetic modifications. In angiosperms, dramatic changes in chromatin state and epigenetic marks take place between dormancy establishment and germination (Lujan-Soto & Dinkova, 2021). Genome-wide or target gene–specific epigenetic analysis should help elucidate the mechanism underlying MpHYP function.

### Possible interacting partners of MpHYP

Mp*HYP* is the sole gene encoding a class VIIIb bHLH transcription factors in *M. polymorpha* (Bowman *et al*., 2017). This class includes Arabidopsis IND and HEC transcription factors, which play critical roles in gynoecium and fruit development (Liljegren *et al*., 2004; Gremski *et al*., 2007; Girin *et al*., 2011; Schuster *et al*., 2015). HEC transcription factors also play important roles in maintaining the shoot apical meristem and in phytochrome-dependent light development (Schuster *et al*., 2014; Zhu *et al*., 2016; Gaillochet *et al*., 2017; Kathare *et al*.,2020). In this study, we demonstrated that MpHYP can promote or repress the expression of its target genes (Figs. 4d and 5b). Such bifunctionality is likely achieved via its dedicated interacting partners. In general, bHLH transcription factors form homodimers or heterodimers with other bHLH proteins. Indeed, IND and HEC interact with other bHLHs such as ALCATRAZ, SPATULA (SPT), and PHYTOCHROME INTERACTING FACTORs (PIFs) (Liljegren *et al*., 2004; Gremski *et al*., 2007; Girin *et al*., 2011; Schuster *et al*., 2014; Zhu *et al*., 2016).

All these proteins belong to the class VII bHLH subfamily, which consists of a single member in *M. polymorpha*, MpPIF (Inoue *et al*., 2016). *Mppif* mutant gemmae show reduced dormancy and germinate even in the dark or inside gemma cups (Inoue *et al*., 2016; Hernandez-Garcia *et al*., 2021), suggesting that MpHYP might function with MpPIF and integrate light and other signals to regulate gemma dormancy. Notably, HECs negatively regulate PIF activity via direct interaction and promote light-dependent seed germination in Arabidopsis (Zhu *et al*., 2016), whereas it appears that MpHYP and MpPIF have an opposite relationship in *M. polymorpha*. The genetic and physical interactions between these bHLH transcription factor genes and their encoding proteins should be carefully investigated in the future. In addition, a recent study indicated that the GRAS-type transcription factor MpDELLA, which is the sole member of the DELLA family in *M. polymorpha*, represses the effect of MpPIF on gemma dormancy (Bowman *et al*., 2017; Hernandez-Garcia *et al*., 2021).

The repression of PIF activity by DELLA via protein–protein interaction is also a conserved mechanism in flowering plants (de Lucas *et al*., 2008; Feng *et al*., 2008). Interestingly, DELLA proteins interact with HEC1 in a yeast two-hybrid assay (Feng *et al*., 2008; Arnaud *et al*., 2010; Gaillochet *et al*., 2018). Identifying the interacting partners of MpHYP would be crucial for understanding the functions and evolution of the class VIIIb bHLH family in land plants.

### Conclusions

Dormancy is important for the survival of land plants and is regulated by the integration of various internal and external cues. The regulatory mechanism of dormancy has been extensively studied in angiosperms, including seeds as well as dormant vegetative organs such as axillary buds and underground bulbs, revealing the significant roles of ABA in these processes (Pan *et al*., 2021). In the present study, we identified MpHYP, a critical regulator of gemma dormancy, in the liverwort *M. polymorpha* and revealed the limited contribution of ABA to the MpHYP-dependent pathway regulating this process. Given the dramatic and specific effect of MpHYP on gemma dormancy, MpHYP might function as a master regulator of this process. Further investigation should provide new insights into the cooperation of ABA-dependent and -independent pathways, the integration of plant hormonal signals and external cues such as light, and the evolution of the functions of class VIIIb bHLH transcription factors in land plants.

## Acknowledgements

We thank Arisa Yasuda for supporting experiments. We appreciate Izumi Yotsui and Yoichi Sakata for providing useful comments on the manuscript. This study was funded by MEXT KAKENHI 17H06472 (K.I.), JSPS KAKENHI Grant Numbers JP19H03247 (K.I.), JP19K23751 and JP21K15125 (H.K.), and JP20K06680 (D.T).

## Author contributions

N.Y., M.Y., H.M., and K.T. conducted the experiments. H.K., N.Y., M.Y., H.M., K.T., D.T., T.F., Y.K., H.F., T.M., and K.I. analyzed the data. H.K. and K.I. designed and supervised the experiments and wrote the article with contributions from all the authors.

## Data availability

RNA-sequence data were deposited at the NCBI in the Sequence Read Archive (SRA) database under accession number DRA013946. The data that support the findings of this study are available from the corresponding author upon reasonable request.

## Supporting information

**Fig. S1.** Knock-out strategy of *MpHYP* by homologous recombination.

**Fig. S2** Mutagenesis of *MpNCED* by CRISPR/Cas9-mediated gene editing.

**Fig. S3** Generation of *Mppyl1^ge^ MpHYP-GR* by CRISPR/Cas9-mediated gene editing.

**Table S1**. Primers used in this study.

**Table S2.** Results of RNA-seq of cell cycle–related genes.

**Table S3**. Results of RNA-seq of ethylene-related genes.

**Table S4.** Results of RNA-seq of auxin-related genes.

**Table S5.** Results of RNA-seq of ABA-related genes.

**Table S6.** Results of RNA-seq of *MpLEAL* genes.

