## Supplementary figures and images for "The bHLH transcription factor MpHYPNOS regulates gemma dormancy in the liverwort *Marchantia polymorpha*"

### Fig. S1

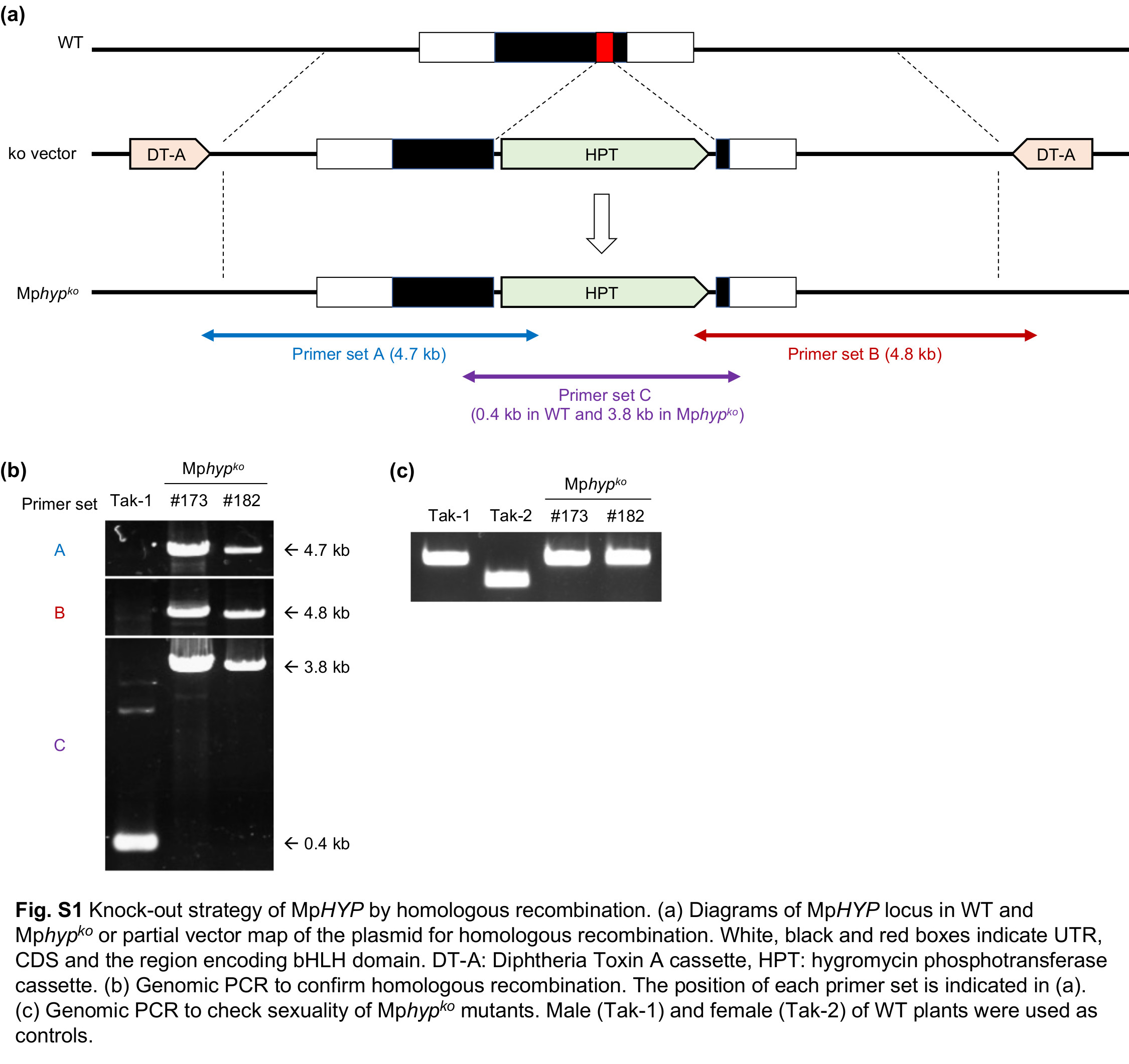

### Fig. S2

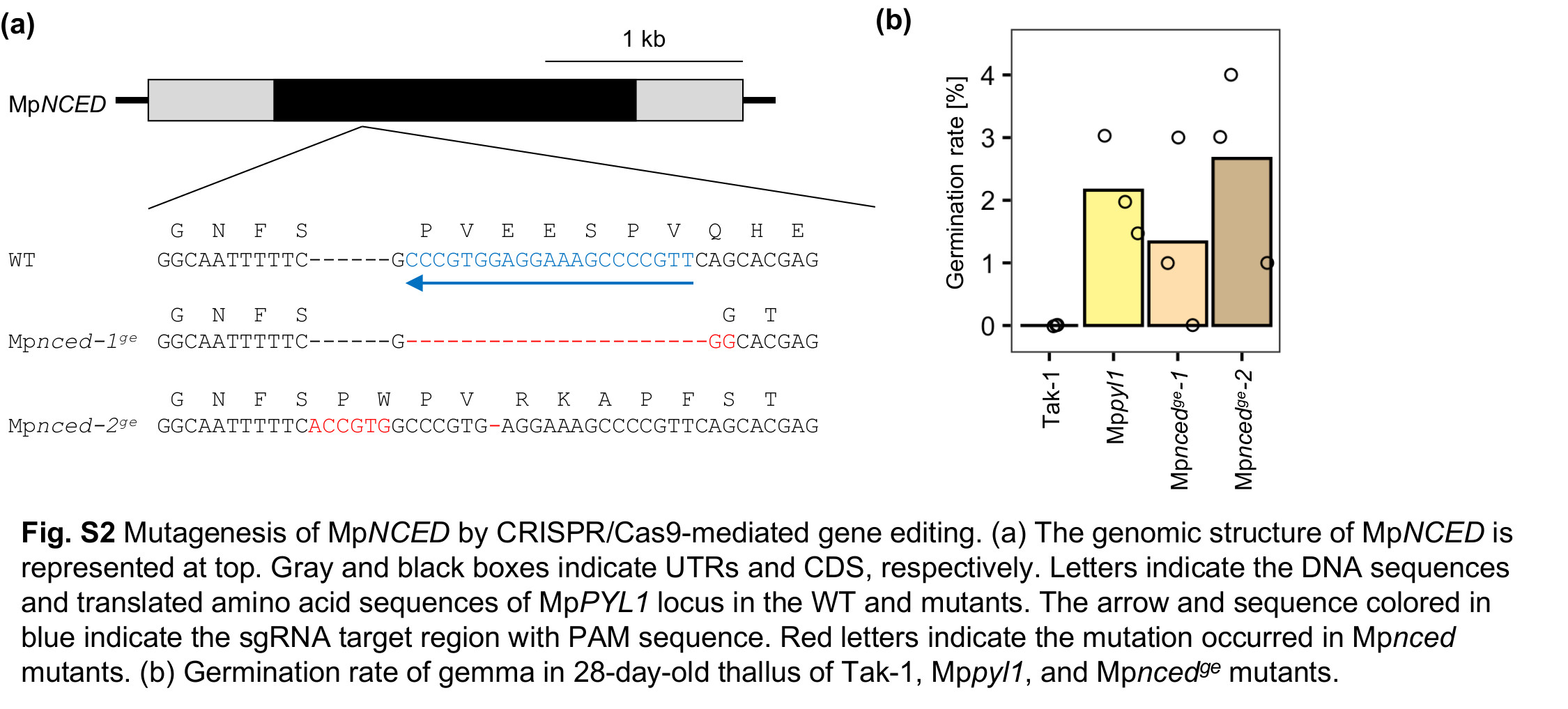

### Fig. S3

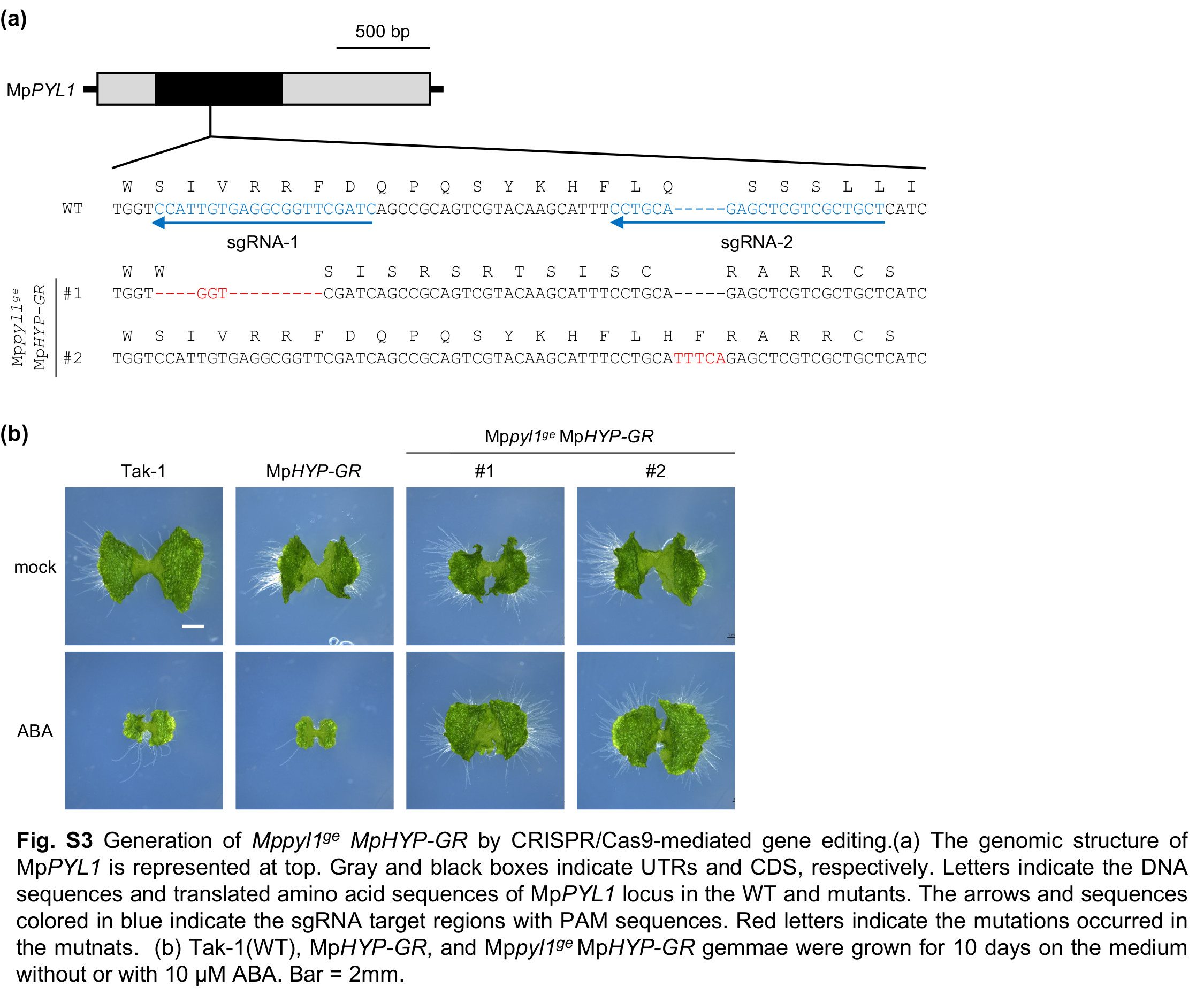
